# Characterizing the Role of Endocarp *A* and *B* Cells Layers During Pod (silique) Development in Brassicaceae

**DOI:** 10.1101/2024.05.21.595188

**Authors:** Justin B. Nichol, Marcus A. Samuel

**Affiliations:** Department of Biological Sciences, University of Calgary, Calgary, AB, Canada T2N 1N4

## Abstract

The process of silique dehiscence is essential for proper dispersal of seeds at the end of a plant’s lifecycle. Current research focuses on genetic manipulation to mitigate this process and enhance shatter tolerance in crop plants, which has significant economic implications. In this study, we have conducted a time-course analysis of cell patterning and development in valve tissues of *Arabidopsis thaliana* and closely related Triangle of U species from Brassicaceae. The goal was to decipher the detailed temporal developmental patterns of the endocarp a and b cell layers of the valve, specifically their degradation and lignification respectively. Additionally, we propose a new classification system for the lignification of the endocarp a cell layer: L1 indicates the cell closest to the replum, with L2 and L3 representing the second and third cells, respectively, each numerical increment indicating lignified cells farther from the replum. Our findings provide a foundational framework absent in current literature, serving as an effective blueprint for future genomic work aimed at modifying valve structures to enhance agronomic traits, such as reducing fiber (lignin) or increasing shatter tolerance.

## Introduction

Classically, *Arabidopsis thaliana* silique development has been described through a flower/carpel developmental staging process that describes morphological markers which define the beginning of each stage (Smyth et al., 1990; Ferrándiz et al., 1999). Using this classical method of flower/carpel development, many nuances are lost in silique development since multiple days are combined into a singular stage (Smyth et al., 1990; Ferrándiz et al., 1999). Taking this into consideration, we developed a silique developmental staging method for *A. thaliana* that is based on days after pollination (DAP), where each cellular developmental change can be associated with a singular time point, helping to capture previously overlooked stages using the classical method. Additionally, we wanted to expand this understanding by applying the principles learned from *Arabidopsis thaliana* to *Brassica napus* (canola) and Triangle of U species as little to no foundational characterization of cell patterning and differentiation during pod (silique) development has been conducted in this area (Spence et al., 1999; Squires et al., 2003; Chen et al., 2011.). Given that canola is an important oil-seed crop in Canada, understanding and characterizing these pods is important for genetic manipulation for advancing trait improvement.

To further understand the silique developmental processes, a thorough understanding of the cellular makeup of the silique is essential. During the reproductive process, following successful fertilization, the gynoecium elongates and undergoes regulated developmental processes to form a mature silique which can be split into two distinct components, the valve and replum (Figure 1a, 3a) (Spence et al. 1996; Dinneny et al. 2005). The valves are two crescent-shaped cell tissues that fuse jointly at a central seam, the replum, with the replum running in parallel down the length of the silique (Figure 1a, 3a-b) (Spence et al. 1996; Dinneny et al. 2005). These two valves are made up of three different types of cells that form the exocarp, mesocarp, and endocarp together forming the pericarp of the silique (Figure 1a-b, Figure 3b-c) (Alvarez et al., 2002; Liljergen et al., 2004; Roeder and Yanofsky, 2006). The exocarp is the outermost layer which comprises the epidermis, a protective barrier for the silique (Figure 1a-b, Figure 3b-c) (Liljergen et al., 2004; Roeder and Yanofsky, 2006). The middle layer, mesocarp, is made primarily of several rows of parenchymal cells making up the majority of the valves (Figure 1a-b, Figure 3b-c) (Liljergen et al., 2004; Roeder and Yanofsky, 2006). The innermost layer comprises the endocarp cells, divided into two distinct cell types, endocarp *a* and *b*, with the endocarp *a* layer being highly vacuolated and the endocarp *b* layer being lignified (Figure 1a-b, Figure 3b-c) (Liljergen et al., 2004; Roeder and Yanofsky, 2006). The replum is the connection point between the two valves on either side (Figure 1a-b, Figure 3b-c) (Liljergen et al., 2004; Roeder and Yanofsky, 2006). The replum forms the main vascular structure of the silique along with the septum, that is connected to it from both ends (Figure 1a-b, Figure 3b-c) (Liljergen et al., 2004; Roeder and Yanofsky, 2006). The fused interaction between the valves and the replum gives rise to the vale margin where a lignified layer of cells connects to the separation layer forming the dehiscence zone (DZ) (Figure 1a-b, Figure 3b-c) (Liljergen et al., 2004; Roeder and Yanofsky, 2006). During the maturation process of the pod, the valve margin begins to separate across the separation layer due to applied tensile forces and water loss (Squires et al., 2003; Liljergen et al., 2004). In conjunction with the hydrolytic enzymes that aid in the process of cell separation, the tensile forces imparted on the lignified cells within the lignified valve layer and the endocarp b (enb) cell layer, from the drying of the mesocarp, lead to separation of the two valves resulting in silique opening (pod shattering) (Squires et al., 2003; Ferrándiz, 2002; Liljergen et al., 2004). During pod opening, the valve and replum split along the DZ allowing the shattering process and the accompanying seed dispersal to occur (Spence et al., 1996; Squires et al., 2003; Meakin an Roberts, 1990; Ferrándiz, 2002; Ferrándiz, 2000).

**Figure 1.**
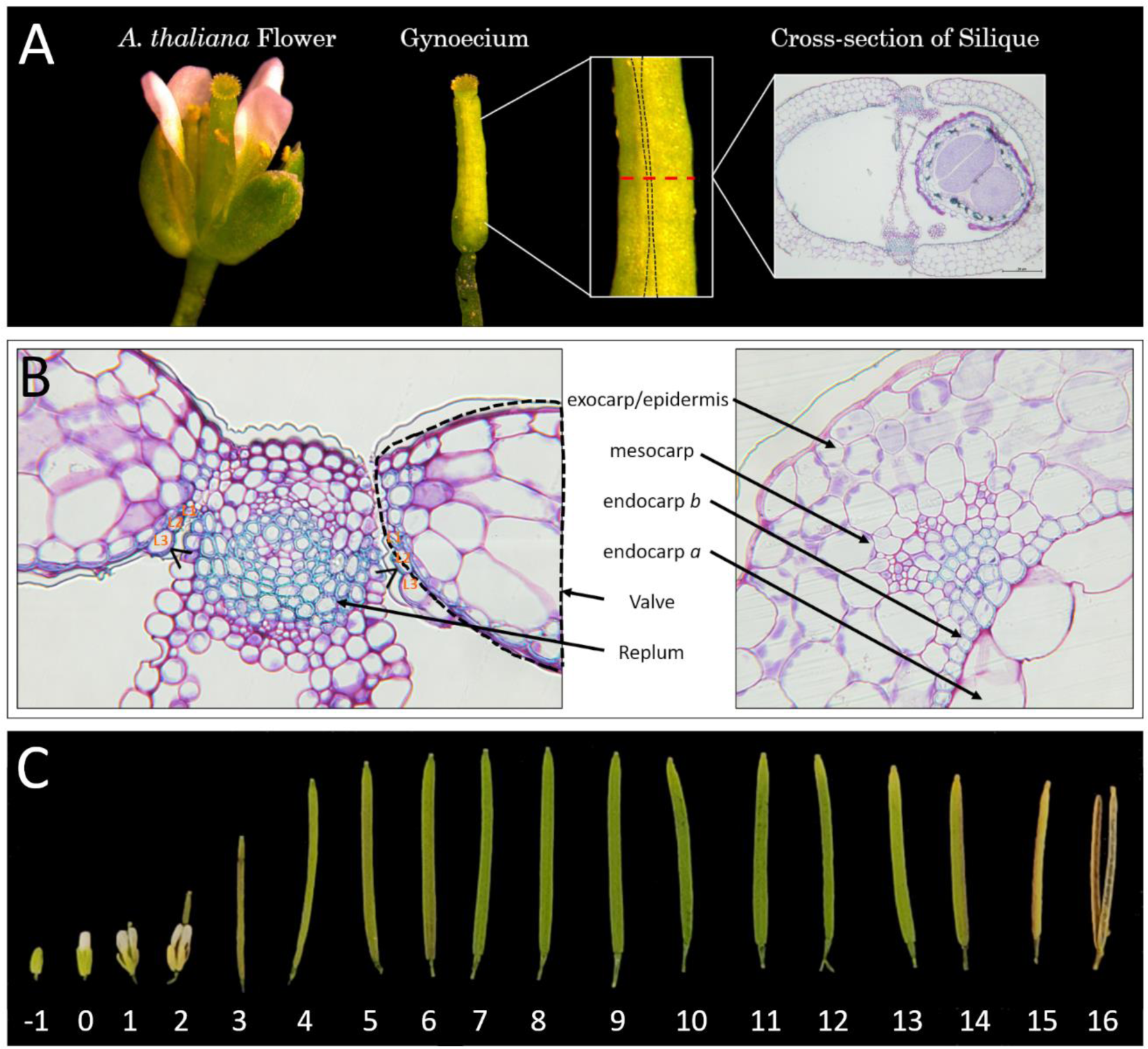
*Arabidopsis thaliana* silique development. A schematic of *Arabidopsis thaliana* ecotype Col-0 cross-sectional collection. The black dotted line representing the replum region and the red dotted line depicting the mid region of the silique where cross-sections were collected with an accompanying representative cross-sectional image (A). The replum and valve region of a wild-type silique cross-section showcasing different cell layers. The black dotted line indicates the valve. Black arrows indicate the endocarp *a* lignification. Endocarp *a* L1-3 cells are indicated on both sides of the replum (B). Whole silique profiling from -1 to 16 days after pollination (DAP)(C).

The goal of this study was to develop a time-course characterization of the cellular development within the silique of *Arabidopsis thaliana* and closely related Triangle of U species. This work aids in laying down the foundation in silique cellular development for future research.

## Materials and methods

### 2.1 Growth conditions and silique date tagging

*Arabidopsis thaliana* plants were grown under standard chamber growth conditions at 22°C with 16 h light and 8 h dark. *Brassica napus* plants were grown under standard greenhouse growth conditions at 22°C ± 4°C with supplemental lights to maintain 16 h light and 8 h dark during winter 2024 (Jan – April 2024). Arabidopsis thaliana Columbia (Col-0) ecotype was used as wild-type. *Brassica napus* (canola) cv. Westar was used as wild-type.

*Arabidopsis thaliana* flowers that had just begun to open were date-tagged each day for 16 days. Flowers for the *Brassica* species were hand-pollinated and tagged each day for 36 days consecutively. Collection occurred for both species one day past the final day.

### 2.2 Sectioning, histochemical staining and whole-mount preparation

*Arabidopsis thaliana* siliques were placed within a general fixative solution of 1.6% paraformaldehyde and 2.5% glutaraldehyde within a 0.05M phosphate buffer at pH 6.8 (Yeung et al., 1996). Fixation of the siliques lasted 24 hrs until a 30-minute vacuum infiltration at 25 mmHg. Siliques were subsequently taken through a graded ethanol dehydration series (50%, 75%, 100%) until an additional vacuum infiltration step. After vacuum infiltration, siliques were placed within a graded HEMA-based resin embedding system (Technovit 7100) solution series with ethanol (50%, 75%) until 100% embedding was achieved (Ruzin 1999). Double-edged razor blades were used to cut embedded siliques to roughly 1mm in size within the mid-region to allow for consistent sectioning results (Figure 1a.). Sections were placed within casts in the desired orientation before the hardening catalyst was added. A Reichert-Jung Autocut 2040 microtome was used to section tissue to 3 µm in size. Sections were placed on to microscope slides to dry in to place. Sections were stained for 1 min using 0.05% Toluidine Blue O (TBO) in 0.05M Citrate Buffer pH 4 (O’Brien et al., 1964). Coverslips were added to the stained section and microscope slide where they were subsequently imaged using a Nixon Ds-Fi2 camera on an Leitz Aristoplan microscope.

*Brassica* species siliques were free hand-sectioned using double-edged razor blades around the mid region to allow for consistency of sections (Figure 1b.) (Ruzin 1999). Sections were stained using the aforementioned TBO protocol in section 2.2. Once sections were stained, they were placed on microscope slides with a coverslip on top. Subsequent images were taken on a Nikon eclipse E200 microscope using a mounted Samsung S23+ Camera.

### 2.3 Maceration preparation

Maceration fluid was prepared by combining one part 30% solution of hydrogen peroxide with four parts distilled water and five parts of glacial acetic acid (Tardif and Conciatori, 2015). The maceration fluid was freshly prepared within a clean glass bottle in the fume hood (Tardif and Conciatori, 2015). *Brassica napus* siliques were cut to 5mm in length and placed within the maceration fluid for 1 to 4 days at 56 degrees Celsius (Tardif and Conciatori, 2015). Once macerated siliques turned whitish in colour, they were washed gently with three changes of water with an hour between each change (Tardif and Conciatori, 2015). Cells of the valve were separated by vigorously shaking them within a vial. Cell mixture was placed on to a glass slide with a coverslip and subsequently imaged using a Nixon Ds-Fi2 camera on an Leitz Aristoplan microscope.

## Results and Discussion

### Endocarp *A* and *B* cellular developmental and lignification patterning in *Arabidopsis thaliana*

To aid in our understanding of silique developmental patterning in *Arabidopsis thaliana*, a complete time course of development was used to understand the cellular patterning of silique development (Figure 1a). Cross-sections were analyzed for each time point of silique development focusing on the different cell layers mentioned in Figure 1b with a complete visualization of silique development from -1 to 16 DAP depicted in Figure 1c. At 3 DAP the siliques lose the other floral organs (Figure 1c). At 6 DAP the silique reaches its maximum length (Figure 1c). The desiccation process of the silique begins near the tip at 11-12 DAP and continues down until it is fully dried at 14-15 DAP (Figure 1c).

Siliques at each individual time point were sectioned from 0 to 16 DAP with the primary focus on the whole cross-section, along with the replum and valve regions (Supplementary Fig 1-5) Silique cross-sections at 3, 6, 9, and 12 DAP were used as representative time points (Figure 2). At 5-6 DAP lignification of the replum and endocarp *b* cell layer within the wild-type *Arabidopsis thaliana* silique appears, with lignification reaching its maximum at 9-10 DAP (Figure 2). It should be noted that endocarp *b* lignification begins in the middle of the valve region at 6 DAP, and over the course of the next 3 days, lignification continues until it reaches the replum at 8-9 DAP (Figure 2) 8 DAP marks the first instance of lignification appearing within the walls of the En*A* 3-Cells until it reaches a maximum at 10 DAP (Figure 2).

**Figure 2.**
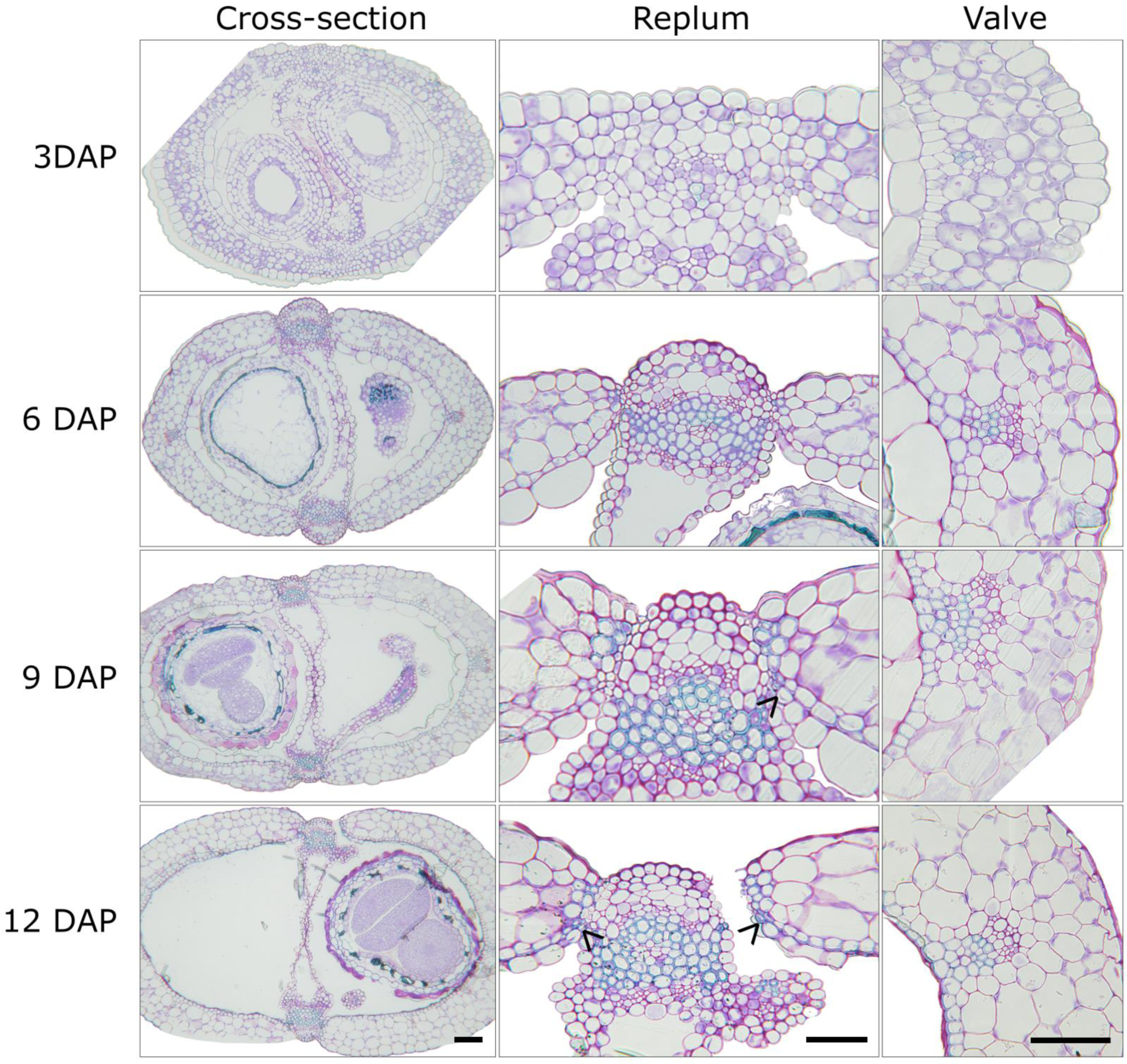
*Arabidopsis thaliana* wild-type silique cross-sections. Wild-type *Arabidopsis thaliana* ecotype Col-0 at 3, 6, 9, 12 days after pollination (DAP) depicting whole silique cross-sections, replum and valve regions. Black arrows indicate the location of endocarp *a* lignification. Scale bars = 50 µm.

The lignification of the endocarp *a* cell layer is a unique process of secondary cell wall deposition since it will only deposit within the walls of the first three cells (Figure 2). Owing to this unique lignin deposition compared to the other cells within the endocarp *a* cell layer, we have decided to classify these cells endocarp *a* L1, L2, and L3 cells. With L1 indicating the closest lignified cell to the replum with L2 and L3 indicating the second and third lignified cell away from the replum, respectively. This new classification system will allow for a more coherent way to describe these cells moving forward. At 9 DAP the endocarp *a* cell layer begins to degrade, with degradation appearing in the middle of the valve region (Figure 2). It is not known whether this degradation is due to PCD or mechanistic drying, although previous research has shown links to GA application and early degradation of endocarp *a* (Spence et al. 1999 and Dorcey et al. 2009). Interestingly, endocarp *a* degradation always appears to follow the lignification of the endocarp *a* L1/2/3 cells, hinting at a potential mechanism between these two processes. The endocarp *a* cell layer finally reaches complete degradation at 12 DAP (Figure 2). The desiccation of the silique begins within the mesocarp layer until it reaches complete desiccation at 15-16 DAP (Figure 1c). Once complete desiccation of the silique has occurred and along with the lignification of the replum and endocarp *b* cell layers, the silique progresses to dehiscence at 16 DAP (Figure 1c). A combination of endocarp *b* lignification and mesocarp desiccation are essential for the precise dehiscence patterning (Spence et al. 1999; Squires et al., 2003; Ferrándiz, 2002; Liljergen et al., 2004; Roeder and Yanofsky, 2006).

### Endocarp *A* and *B* cellular developmental and lignification patterning in *Brassica napus*

Taking our knowledge gathered from *A. thaliana* silique development, we looked to further deepen our understanding of silique (commonly known as pod) developmental patterning in a common oil crop species, canola. *A. thaliana* and canola share many similarities between their flower and gynoecium/silique development since they are closely related species (Figure 1a and 3a) (Zuñiga-Mayo et al., 2018). A representative whole silique cross-section is shown in Figure 3b along with a complete time-course of pod morphology and development from 0 to 35 DAP shown in Figure 3c. At 4 DAP the siliques lose the other floral organs and at 9-10 DAP the silique reaches its maximum length (Figure 3c). Drying of the siliques is shown to begin at 30-31 DAP until desiccation is complete at 35 DAP (Figure 3d).

**Figure 3.**
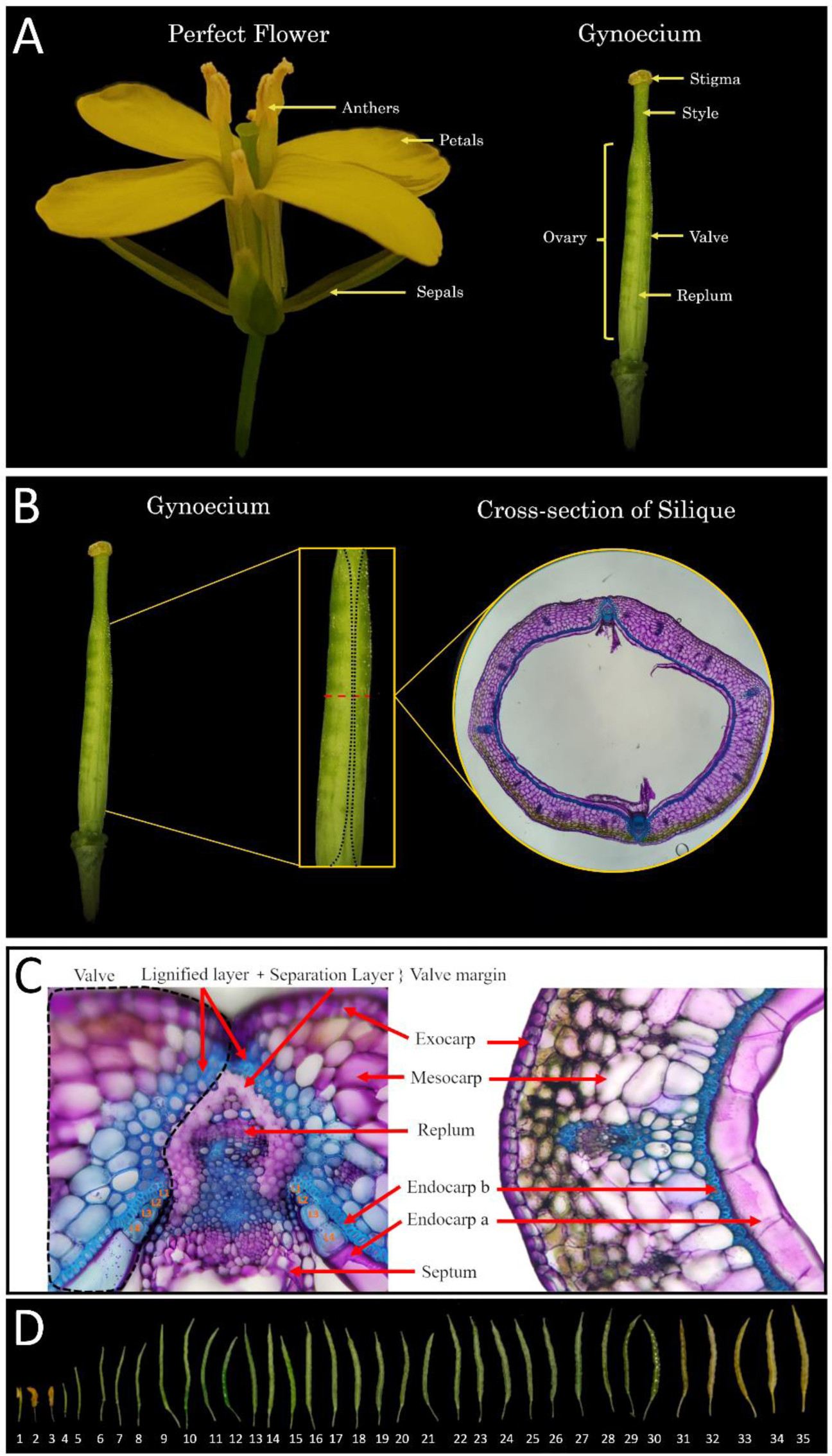
*Brassica napus* silique development. A schematic of *Brassica napus* westar cross-sectional collection. The black dotted line represents the replum region and the red dotted line depicts the mid-region of the silique where cross-sections were collected with an accompanying representative cross-sectional image (A-B). The replum and valve region of a wild-type silique cross-section showcasing different cell layers. The black dotted line indicates the valve. Endocarp *a* L1-4 cells are indicated on both sides of the replum (C). Whole silique profiling from 1 to 35 days after pollination (DAP)(D).

Similar to *Arabidopsis*, canola siliques contain two replum regions on opposite ends of the cross-section along with two valves on either side (Figure 3b-c). Canola valves expand over top of the replum, covering the separation layer (Figure 3c). Lignification can also extend further into the mesocarp layer of the valve as the silique matures (Figure 3c). To further expand our knowledge of the cellular makeup of the endocarp *a* and *b* cell layers, the vales were macerated from mature canola siliques (Figure 4a-c). We found that the lignified endocarp *a* cells along with the lignified mesocarp layer are predominantly thick-walled parenchyma with identifiable plasmodesmata pits lining the exterior walls (Figure 4a-b). In contrast, the endocarp *b* cell layer is comprised of sclerenchyma (Figure 4c). From temporal profiling through TBO staining, it can be observed that lignification in the mesocarp layer begins around 20 DAP following initial lignification of the endocarp *b* cell layer (Figure 4d). The lignification process of the endocarp *b* cell layer begins at 12 DAP with the walls continuing to thicken over the course of several days (Figure 5). It has previously been shown by Jaradat et al. 2014, that the endocarp *a* cell layer undergoes the same lignification in *Brassica* species that is also observed in *A. thaliana*. The variation of the amount of these cells differs between section and species (Jaradat et al. 2014).

**Figure 4.**
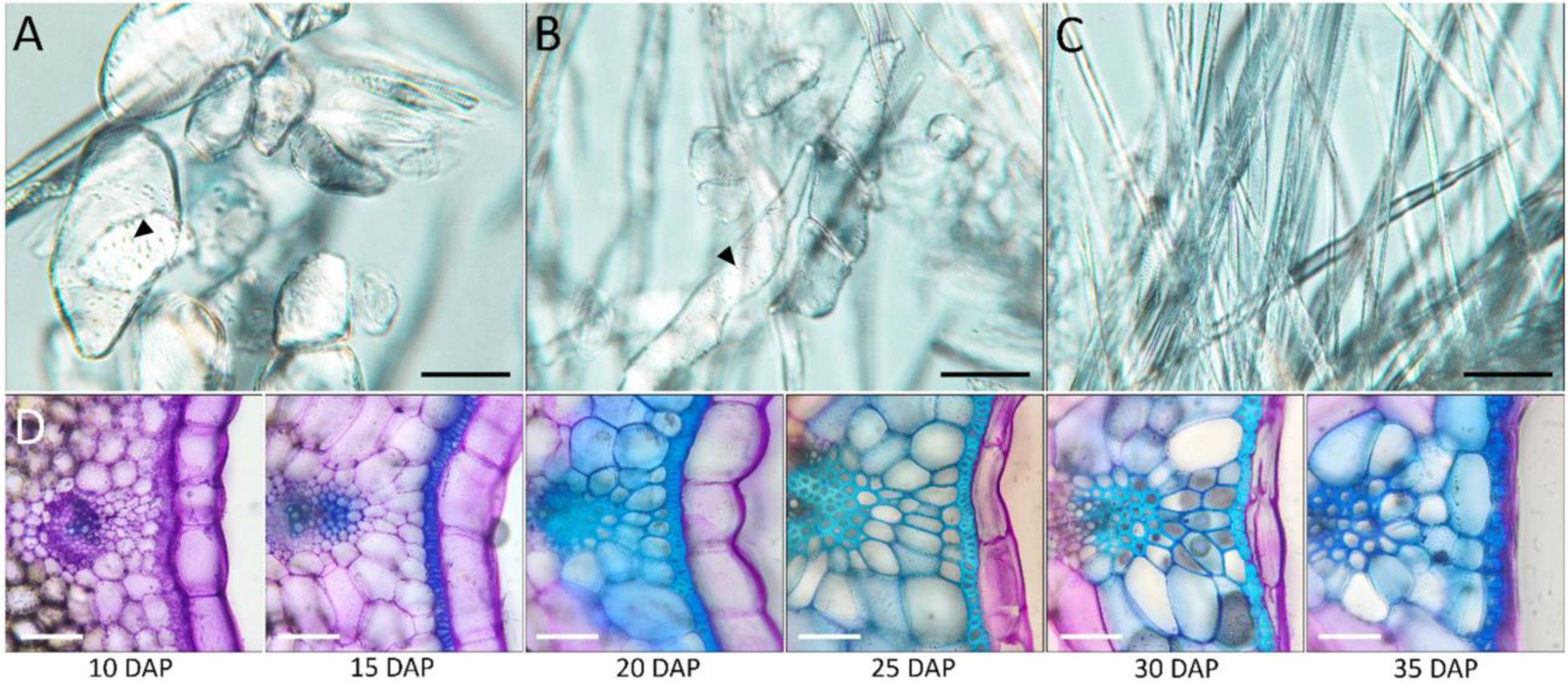
Valve development and cell types. Macerated valve regions of mature *Brassica napus* wild-type Westar siliques depicting thick-walled parenchyma cells. Black arrows indicate plasmodesmata pits (A-B) and sclerenchyma cells (C). Cross-sections of the valve region at 10, 15, 20, 25, 30, 35 days after pollination (DAP) depicting En*B* lignification and En*A* collapse (D). Scale bars = 25 µm.

**Figure 5.**
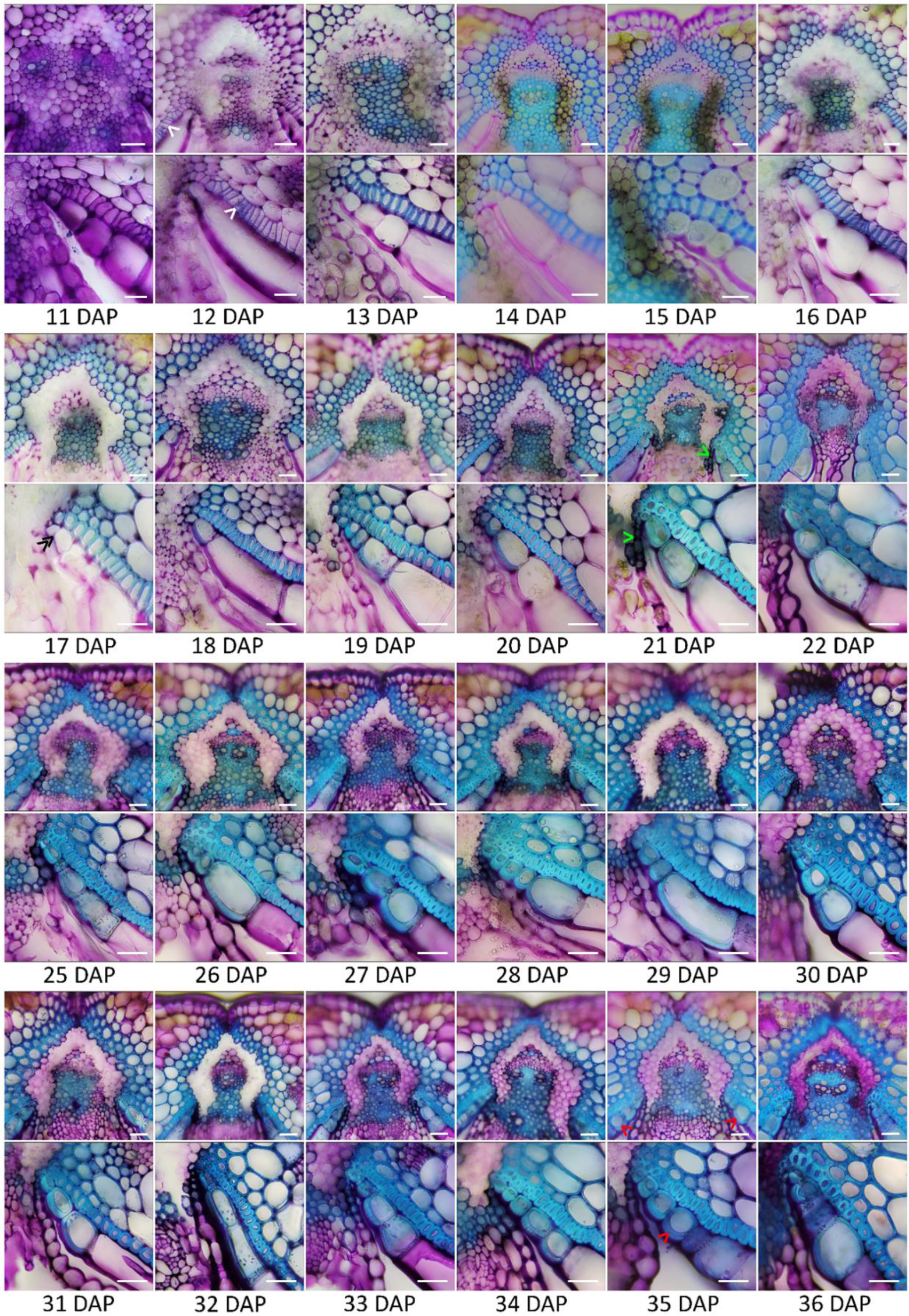
Temporal lignification of *Brassica napus* silique. Wild-type *Brassica napus* Westar siliques taken 11-22, 25-27 days after pollination (DAP) depicting cross-sections of the replum and EnA 3-Cell regions. White arrows indicate location of initial lignification within the endocarp *b* cell layer. Double black arrows indicate the initial lignification of the endocarp *a* cell layer. Green arrows indicate the first instance of the lignification linking the replum and endocarp *a/b* junction. Red arrows indicate the asymmetric thickening of the endocarp *a* L3 cells. Scale bars = 25 µm.

In line with this previous research and the new classification system we have put forth in *A. thaliana*, we decided to acknowledge these unique lignified cells of the endocarp *a* layer in a similar manner with the closest lignified cell to the replum as L1, increasing numerically until the final instance of lignification is seen. At 17 DAP, the first instance of lignification occurring in the endocarp *a* L1 cell is seen (Figure 5). Similar to *Arabidopsis*, the lignification appears in the first cell closest to the replum before working outward (Figure 5). By 18 DAP, the lignification has progressed to the L2 cell and by 20 DAP the lignification has progressed to L3 (Figure 5). Between 33 and 35 DAP the endocarp *a* lignification has progressed to an L4 and L5 cell, with L5 being the last lignified endocarp *a* cell from the replum (Figure 5.). The endocarp *a* L1/2/3 cells continue to thicken after 20 DAP leading to an asymmetric deposition of lignin on the inner wall closest to the ovary space, until a maximum asymmetric deposition of lignin is reached at 35 DAP (Figure 5). Endocarp *a* cells begin to degrade 25 DAP until they are no longer visible at 35 DAP (Figure 4d). Similar to *A. thaliana*, the degradation of the endocarp *a* cell layer follows the lignification of the endocarp *a* L1/2/3 cells (Figure 5). Between 19 and 22 DAP, a small file of lignified cells was seen to connect between the replum and the junction of the *endocarp a* and *b* cell layers (Figure 5).

Degradation of the separation layer is shown to begin at 11 DAP with minimal staining (pectin is stain in pink) of the separation layer remaining after this time point (Figure 5). The lack of colour indicates a complete degradation of the pectic substances within the middle lamella and cell wall respectively (O’Brien et al., 1964; Squires et al., 2003; Meakin and Roberts, 1990; Sander et al., 2001; He et al., 2018; Fry et al., 1992; Jenkins et a., 1999; Petersen et al., 1996; Jenkins et a., 1996). To prime the silique for dehiscence, a combination of the cell wall degradation of the separation layer middle lamella along with the endocarp *b* layer lignification is required (Meakin and Roberts, 1990; Sander et al., 2001; He et al., 2018; Fry et al., 1992; Jenkins et al., 1999; Petersen et al., 1996; Jenkins et a., 1996). As shown in figure 5, the first step in silique dehiscence priming of the middle lamella and cell wall degradation comes much earlier than the complete lignification of the replum and endocarp *a/b* layers.

### Understanding Brassicaceae silique cellular developmental patterning

Expanding our understanding of silique development led us to look into the development of other closely related species to *A. thaliana* and *B. napus* within the Brassicaceae family. We decided to focus on species within the Triangle of U, which includes the six most commonly known members of the genus *Brassica*. Through allopolyploidization events, the three ancestral diploid species combined to create the three common tetraploid crop species (Chen et al., 2011). We investigated silique cellular development and patterning for the 3 tetraploid species *Brassica juncea* (AABB)*, Brassica carinata* (BBCC), and *Brassica napus* (AACC) as well as two of the three diploid species *Brassica rapa* (AA), and *Brassica nigra* (BB). It should be noted that due to difficulties inducing bolting and flowering in *Brassica oleracea* (CC), we could not obtain a time-course of siliques for this species.

Focusing on the similarities and differences between the homeologous species, we observed slight variations in the silique morphology from 0 to 35 DAP (Supplementary Figure 6). *B nigra* had the flattest pod morphology and *B. rapa* had the smallest pod over the time-course amongst the Triangle of U species (Supplementary Figure 6). When looking at the cellular patterning and development, lignification occurs before 10 DAP in *B. rapa* and *B. juncea* compared to later lignification in *B. napus, B. carinata,* and *B. nigra* (Supplementary Figures 7-11). Mesocarp lignification within the valve region is pronounced within *B. nigra* and *B. rapa* siliques compared to the other homeologs (Supplementary Figures 9 and 11). Interestingly, noticeable gaps appear within the endocarp *b* cell layer within *B. napus, B. juncea, B. rapa,* and *B. nigra* but are not found within *B. carinata* (Supplementary Figures 7-11). For all of the Triangle of U species, lignification is seen within the first three cells of the endocarp *a* cell layer (Supplementary Figures 7-11). The size of the endocarp *a* L3-cells differs with *B. juncea, B. nigra,* and *B. rapa* having a reduction in the overall size and shape of the cell (Supplementary Figure 8, 10-11). The largest endocarp *a* L3-cell is found within *B. napus* and *B. rapa* (Supplementary Figure 7 and 9). This data is in agreement to the work done by Jaradat et al. 2014, which also found the size of the endocarp *a* L3-cells in *B. juncea* to be much smaller than *B. napus*.

Endocarp *a* degradation in all species begins at 25 DAP and is seen to be completely degraded by 35 DAP (Supplementary Figures 7-11). Complete degradation is shown to be slightly earlier in *B. juncea* and *B. nigra* (Supplementary Figure 8 and 11). This confirms previous results shown in the other Brassicaceae members where L1/2/3 lignification occurs prior to endocarp a degradation. B*. nigra* has a pronounced bulge near the primary vasculature of the valve beginning at 15 DAP (Supplementary Figure 11). This same phenotype is marginally seen within *B. rapa* and *B. juncea* (Supplementary Figure 8 and 9).

The connection between the phenotypic differences can be better understood when comparing it between the two diploid species and the three tetraploid species. The earlier lignification of *B. rapa* and *B. juncea* is likely due to the A genome since *B. nigra* only comprises of the B genome. The size of the endocarp *a* 3-cells differs within the three tetraploid species of *B. carinata, B. juncea,* and *B. napus* compared to the two diploid species. Since *B. nigra* is observed to have the smallest endocarp *a* 3-cells, this trait is likely passed on through the B genome to *B. carinata* and *B. juncea* which also have relatively small endocarp *a* 3-cells. *B. napus* has the largest endocarp *a* 3-cells most likely due to the C genome in combination with the A genome. The early degradation patterning seen in the endocarp *a* layer in *B. nigra* and *B. juncea* is likely connected to the B genome. All this taken into consideration, future genetics-based research approaches are needed to support these phenotypic claims and confirm the link to the correct genome copy amongst the Triangle of U species. Nevertheless, our descriptive work in this study will serve as a blueprint for the scientific community engaged in genomics work to manipulate valve structure to enhance agronomic traits such as reducing lignification or enhancing shatter tolerance.

## Acknowledgements

We acknowledge NSERC discovery grant for funding for this work. We also thank Shakshi Dutt for her valuable input on edits of this paper.

---

**Supplementary Figure 1.**
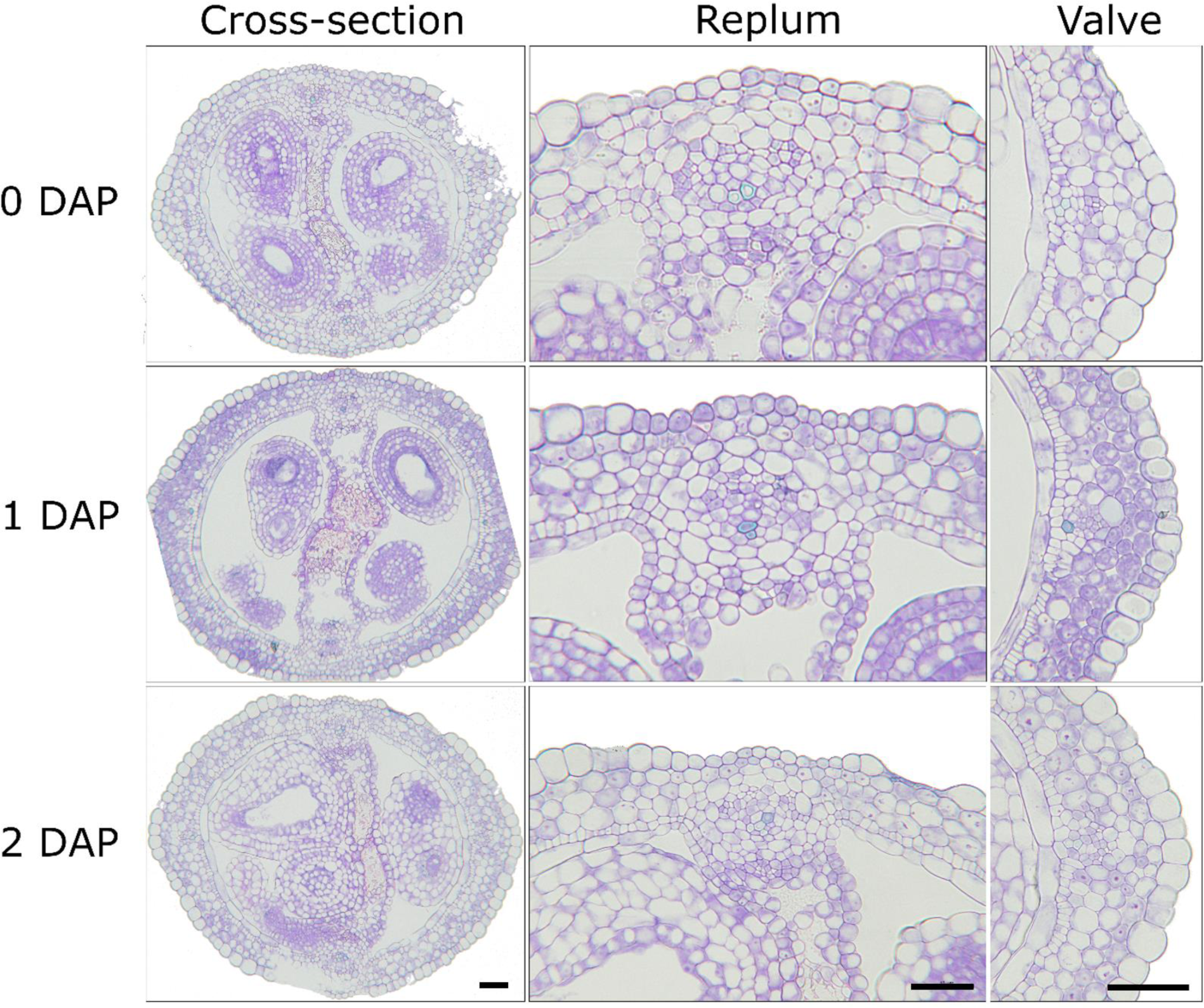
*Arabidopsis thaliana* wild-type silique cross-sections. Wild-type *Arabidopsis thaliana* ecotype Col-0 at 0-2 days after pollination (DAP) depicting whole silique cross-sections, replum and valve regions. Scale bars = 50 µm for each of the images in the panel.

**Supplementary Figure 2.**
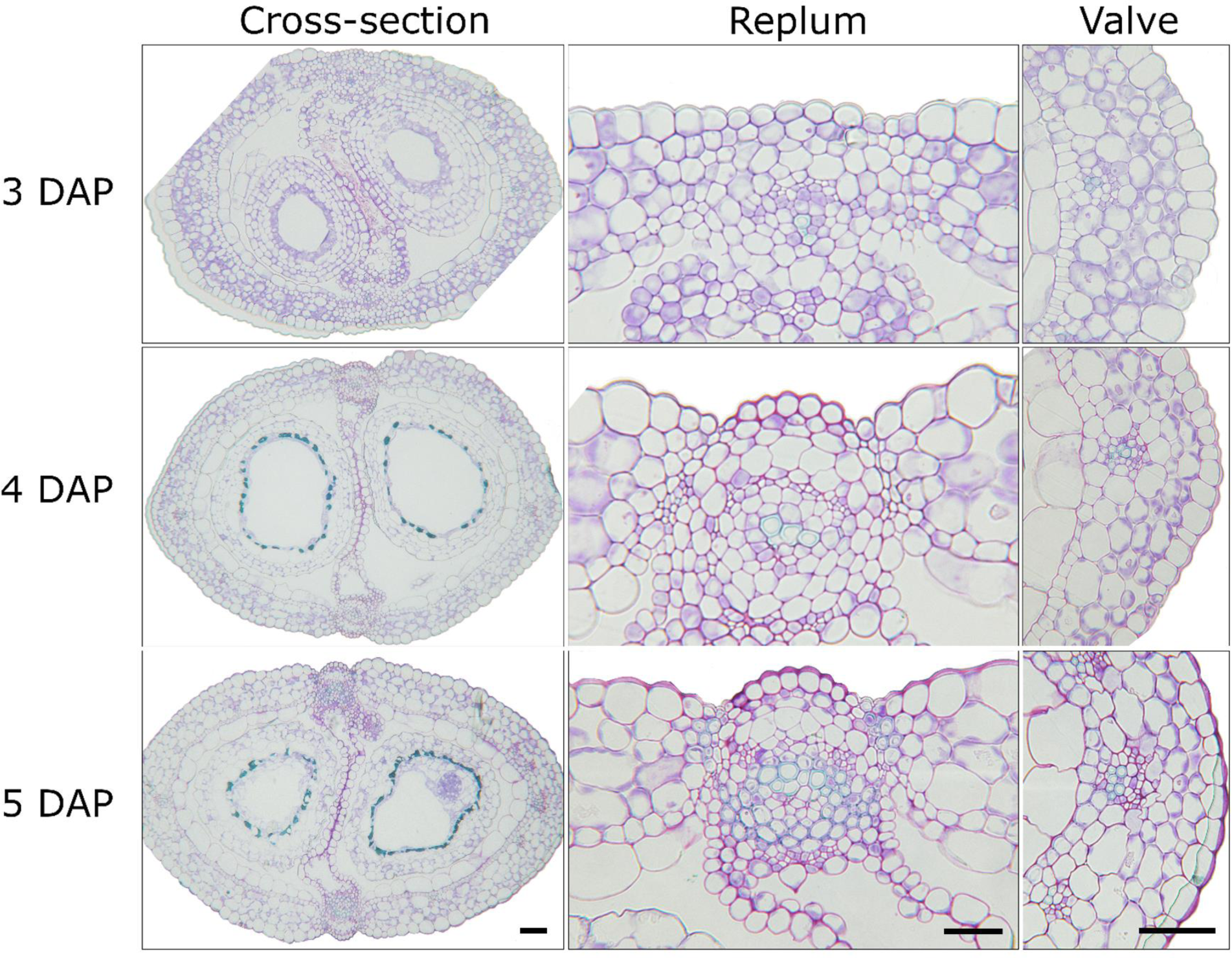
*Arabidopsis thaliana* wild-type silique cross-sections. Wild-type *Arabidopsis thaliana* ecotype Col-0 at 3-5 days after pollination (DAP) depicting whole silique cross-sections, replum and valve regions. Scale bars = 50 µm for each of the images in the panel.

**Supplementary Figure 3.**
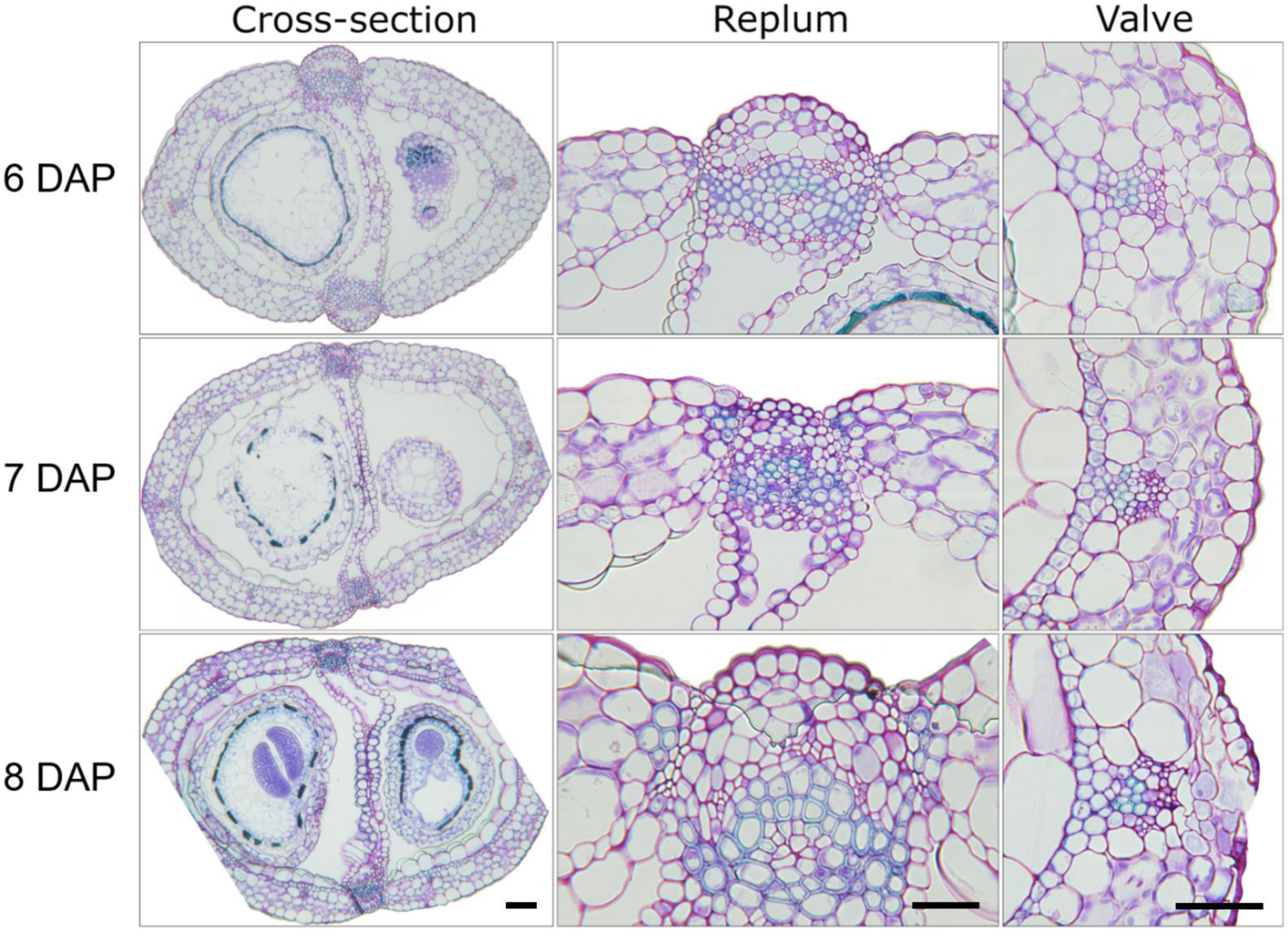
*Arabidopsis thaliana* wild-type silique cross-sections. Wild-type *Arabidopsis thaliana* ecotype Col-0 at 6-8 days after pollination (DAP) depicting whole silique cross-sections, replum and valve regions. Scale bars = 50 µm for each of the images in the panel.

**Supplementary Figure 4.**
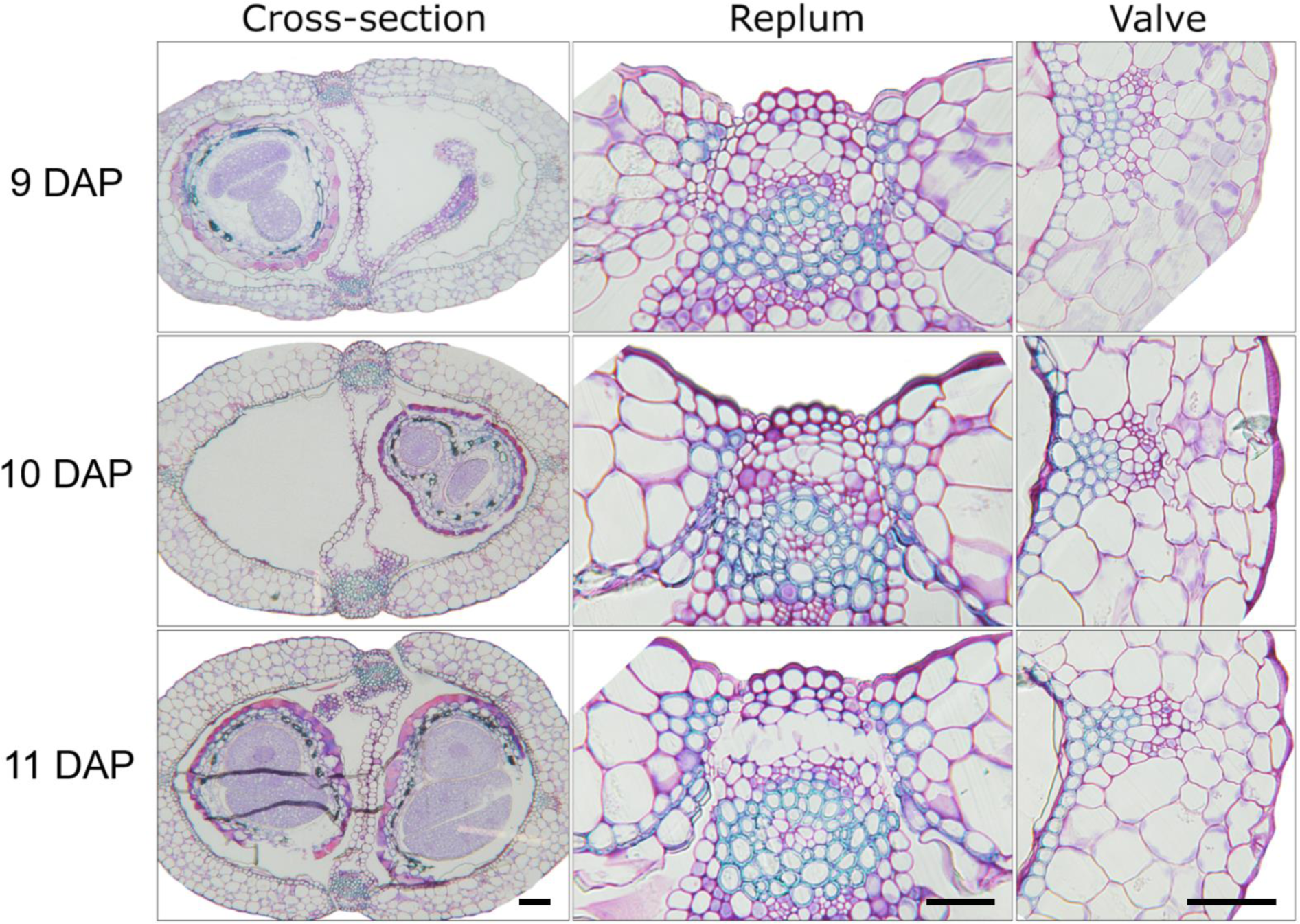
*Arabidopsis thaliana* wild-type silique cross-sections. Wild-type *Arabidopsis thaliana* ecotype Col-0 at 9-11 days after pollination (DAP) depicting whole silique cross-sections, replum and valve regions. Scale bars = 50 µm for each of the images in the panel.

**Supplementary Figure 5.**
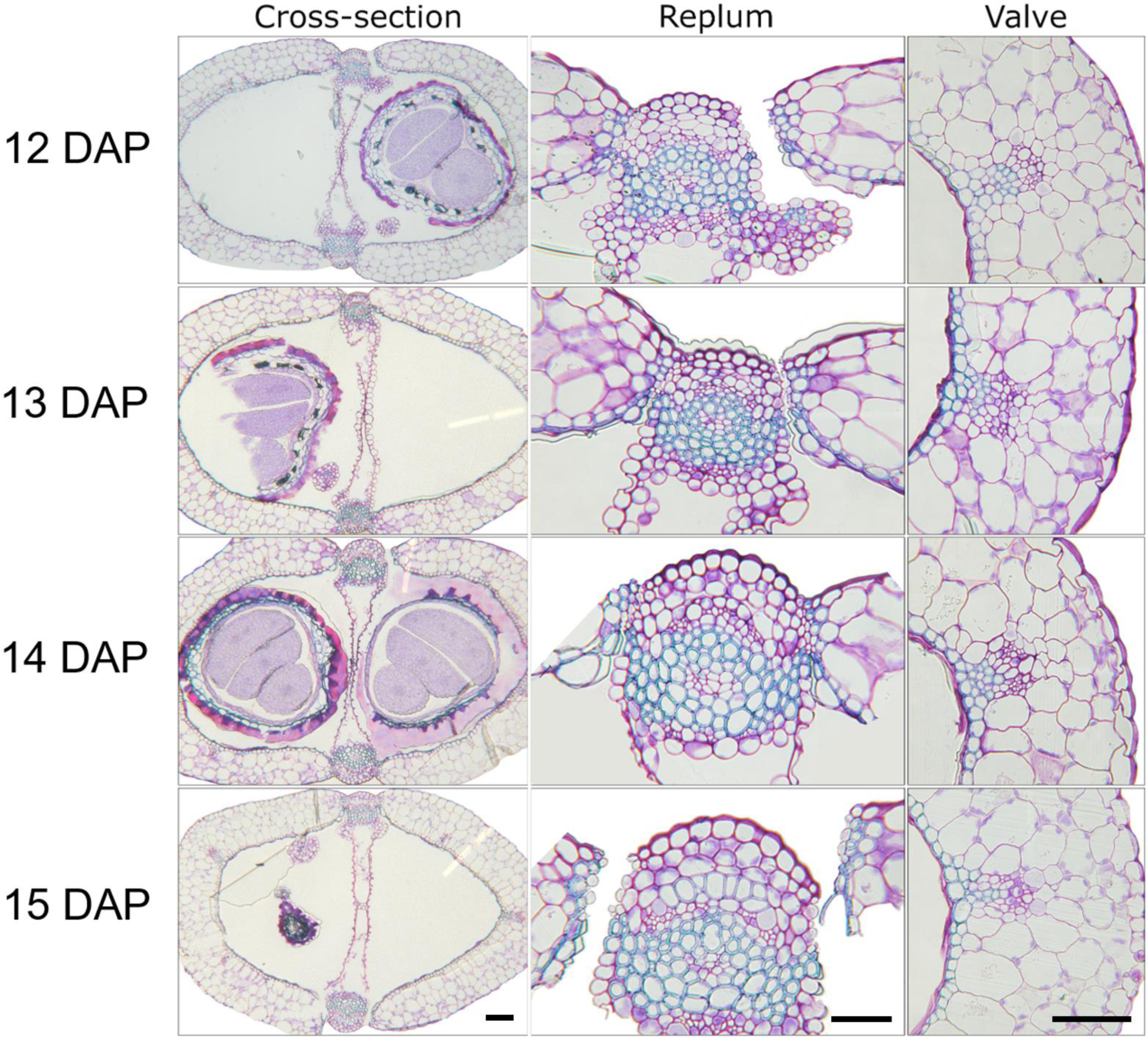
*Arabidopsis thaliana* wild-type silique cross-sections. Wild-type *Arabidopsis thaliana* ecotype Col-0 at 12-16 days after pollination (DAP) depicting whole silique cross-sections, replum and valve regions. Scale bars = 50 µm for each of the images in the panel.

**Supplementary Figure 6.**
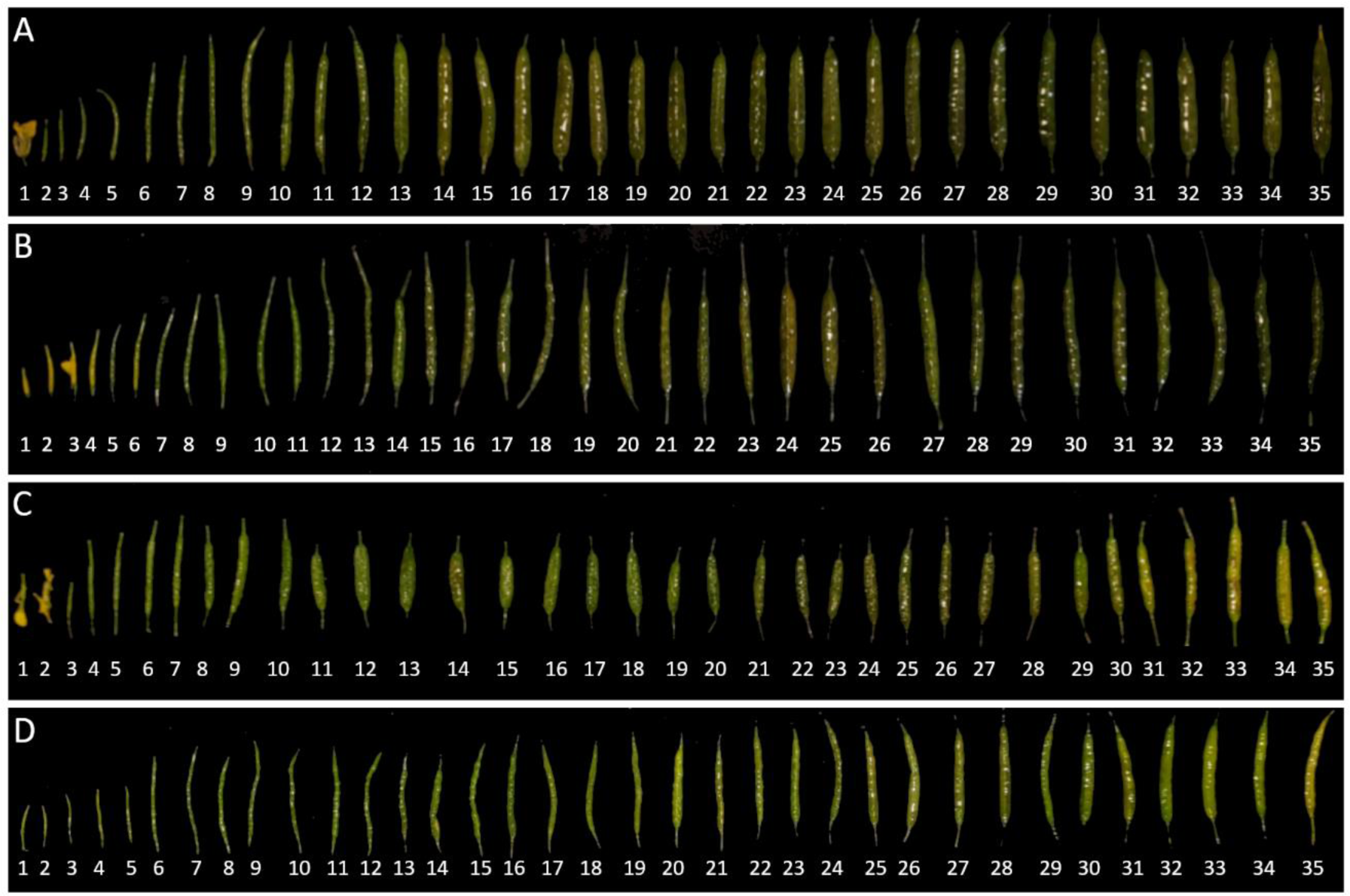
Temporal development of whole Brassicaceae siliques. Brassicaceae whole siliques from 1 to 35 days after pollination (DAP) which include, *Brassica carinata* (A), *Brassica nigra* (B), *Brassica rapa* (C), and *Brassica juncea* (D).

**Supplementary Figure 7.**
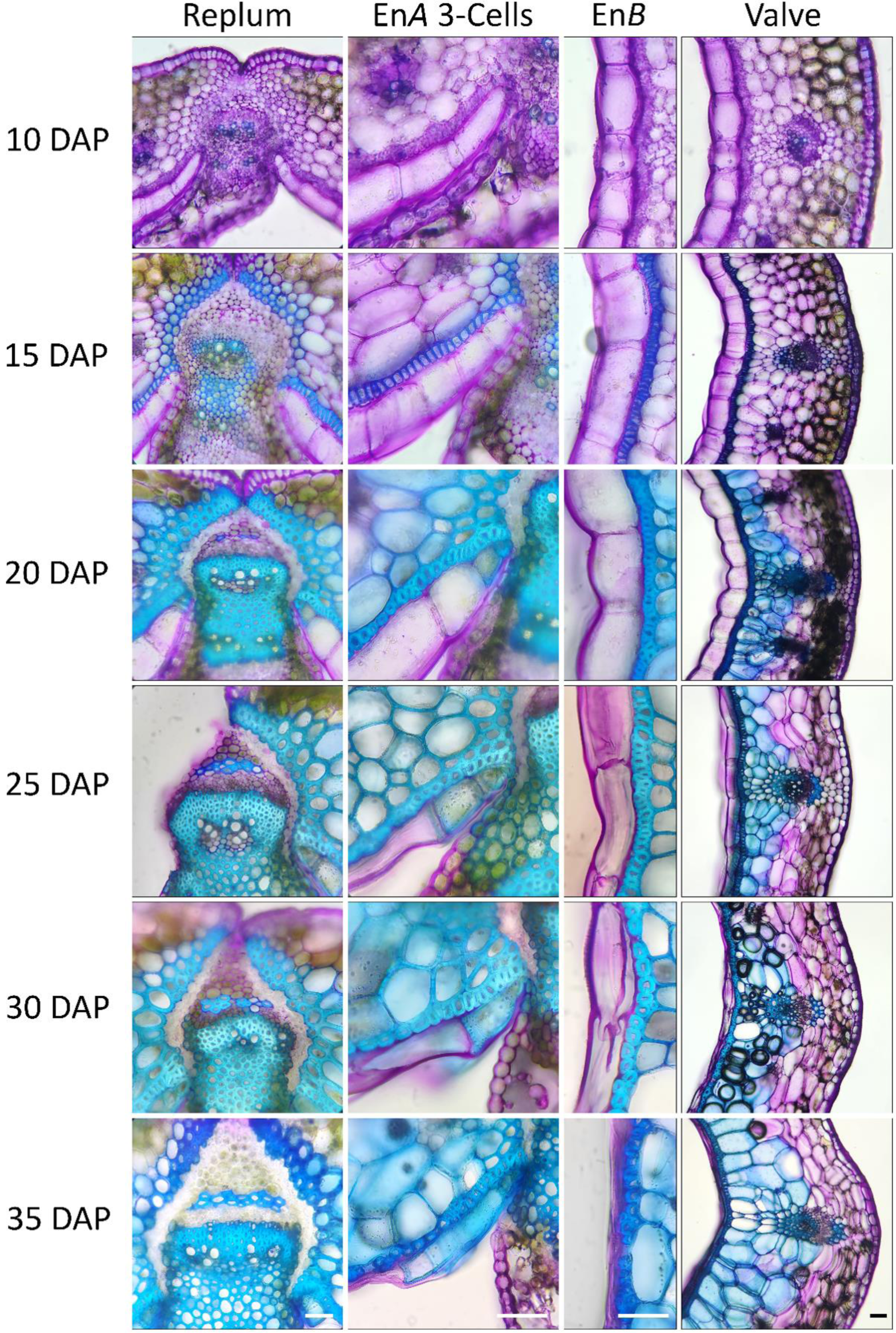
Spatial and temporal lignification of *Brassica napus* siliques. Whole silique cross-sectional images 10, 15, 20, 25, 30, 35 days after pollination (DAP) depicting the replum, En*A* 3-Cells, En*B*, and valve spatial and temporal lignification patterning. Scale bars = 25 µm for each of the images in the panel.

**Supplementary Figure 8.**
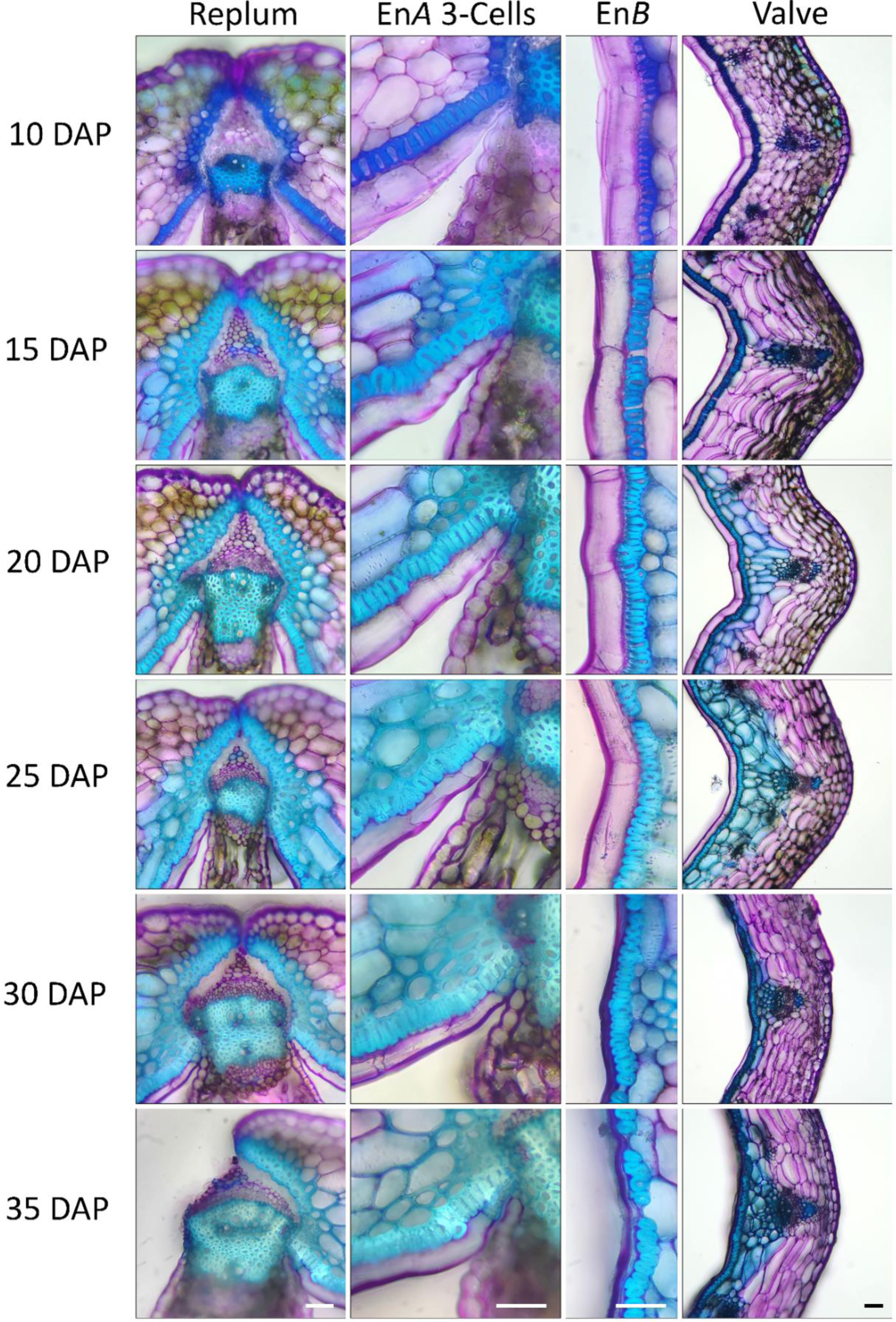
Spatial and temporal lignification of *Brassica juncea* siliques. Whole silique cross-sectional images 10, 15, 20, 25, 30, 35 days after pollination (DAP) depicting the replum, En*A* 3-Cells, En*B*, and valve spatial and temporal lignification patterning. Scale bars = 25 µm for each of the images in the panel.

**Supplementary Figure 9.**
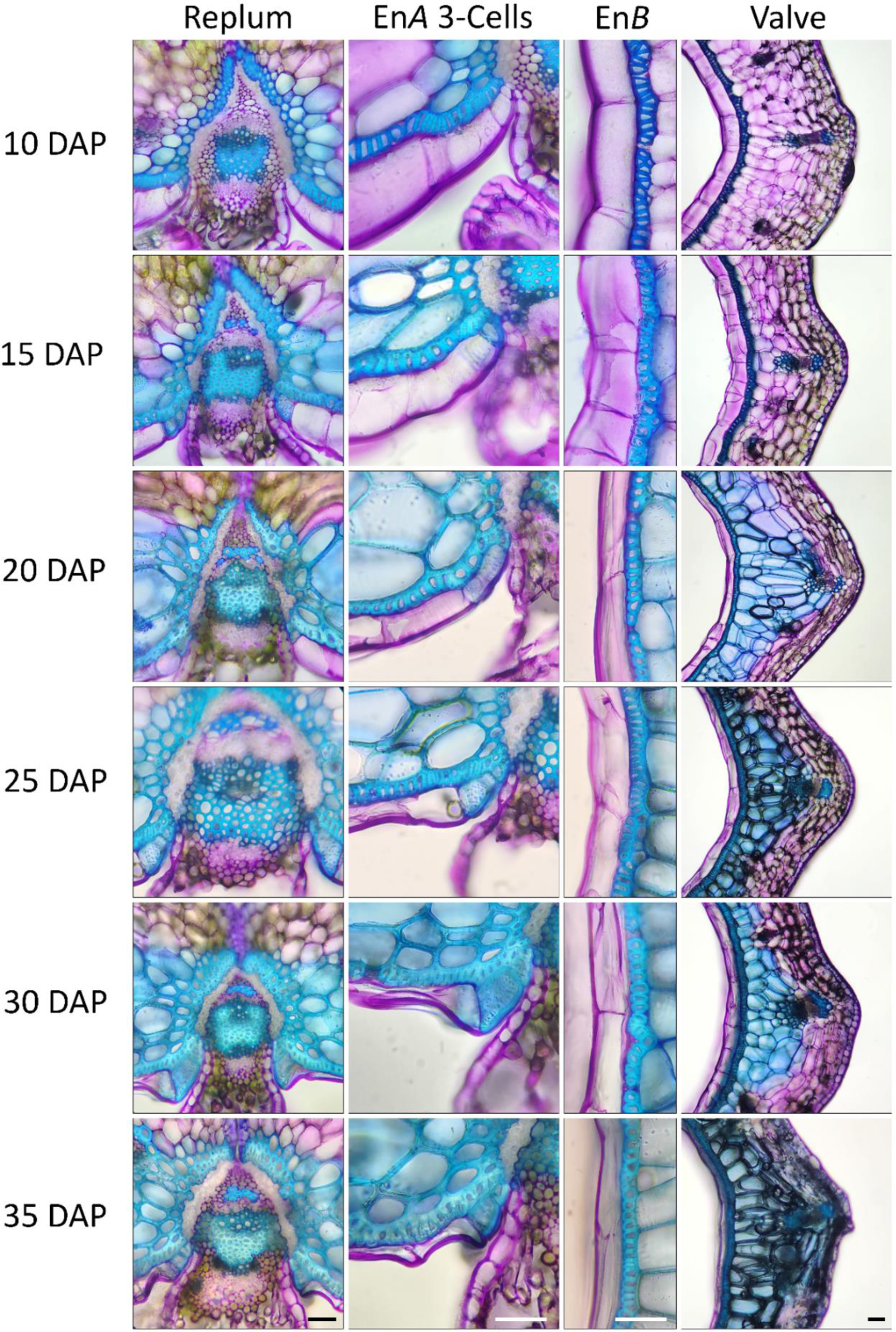
Spatial and temporal lignification of *Brassica rapa* siliques. Whole silique cross-sectional images 10, 15, 20, 25, 30, 35 days after pollination (DAP) depicting the replum, En*A* 3-Cells, En*B*, and valve spatial and temporal lignification patterning. Scale bars = 25 µm for each of the images in the panel.

**Supplementary Figure 10.**
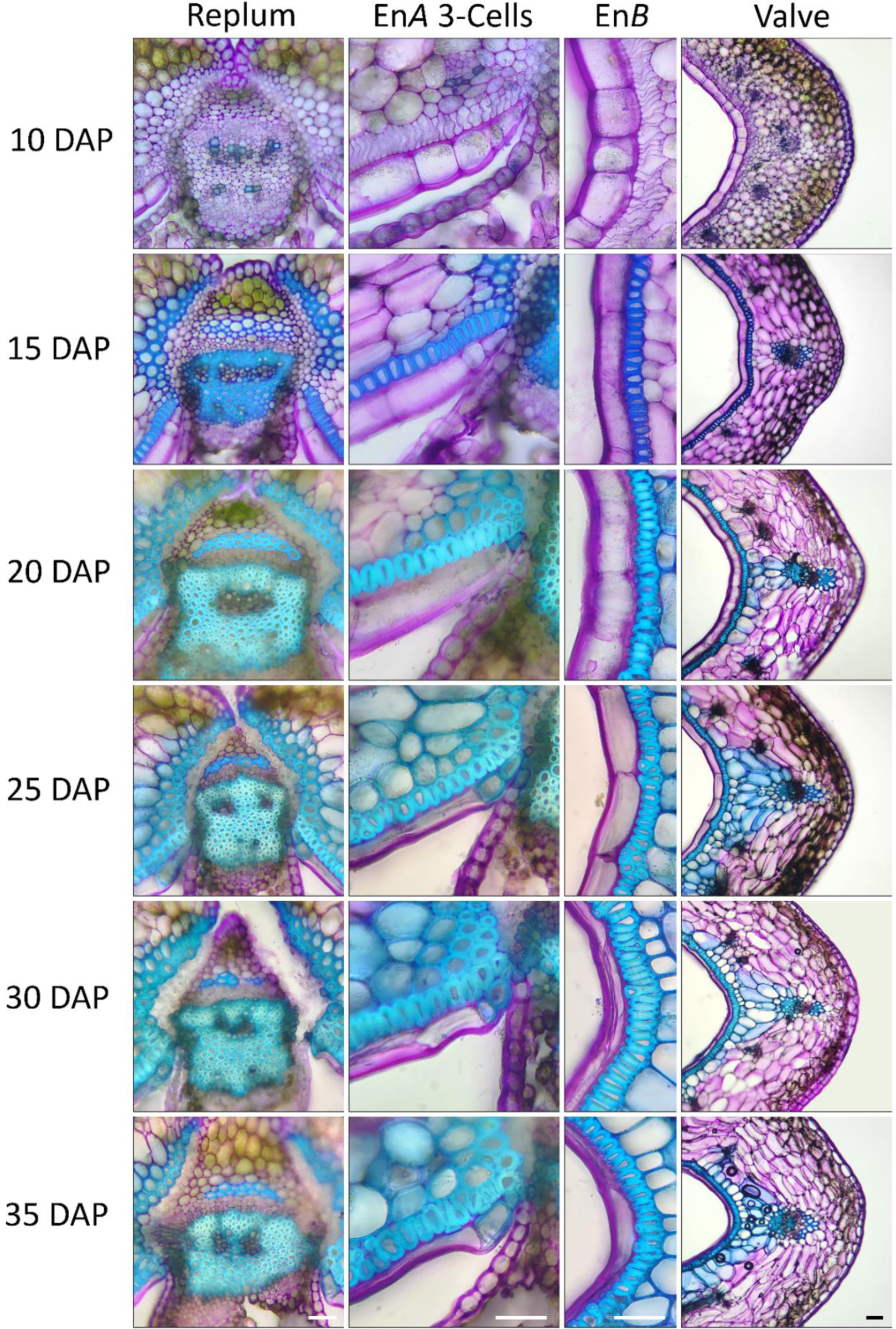
Spatial and temporal lignification of *Brassica carinata* siliques. Whole silique cross-sectional images 10, 15, 20, 25, 30, 35 days after pollination (DAP) depicting the replum, En*A* 3-Cells, En*B*, and valve spatial and temporal lignification patterning. Scale bars = 25 µm for each of the images in the panel.

**Supplementary Figure 11.**
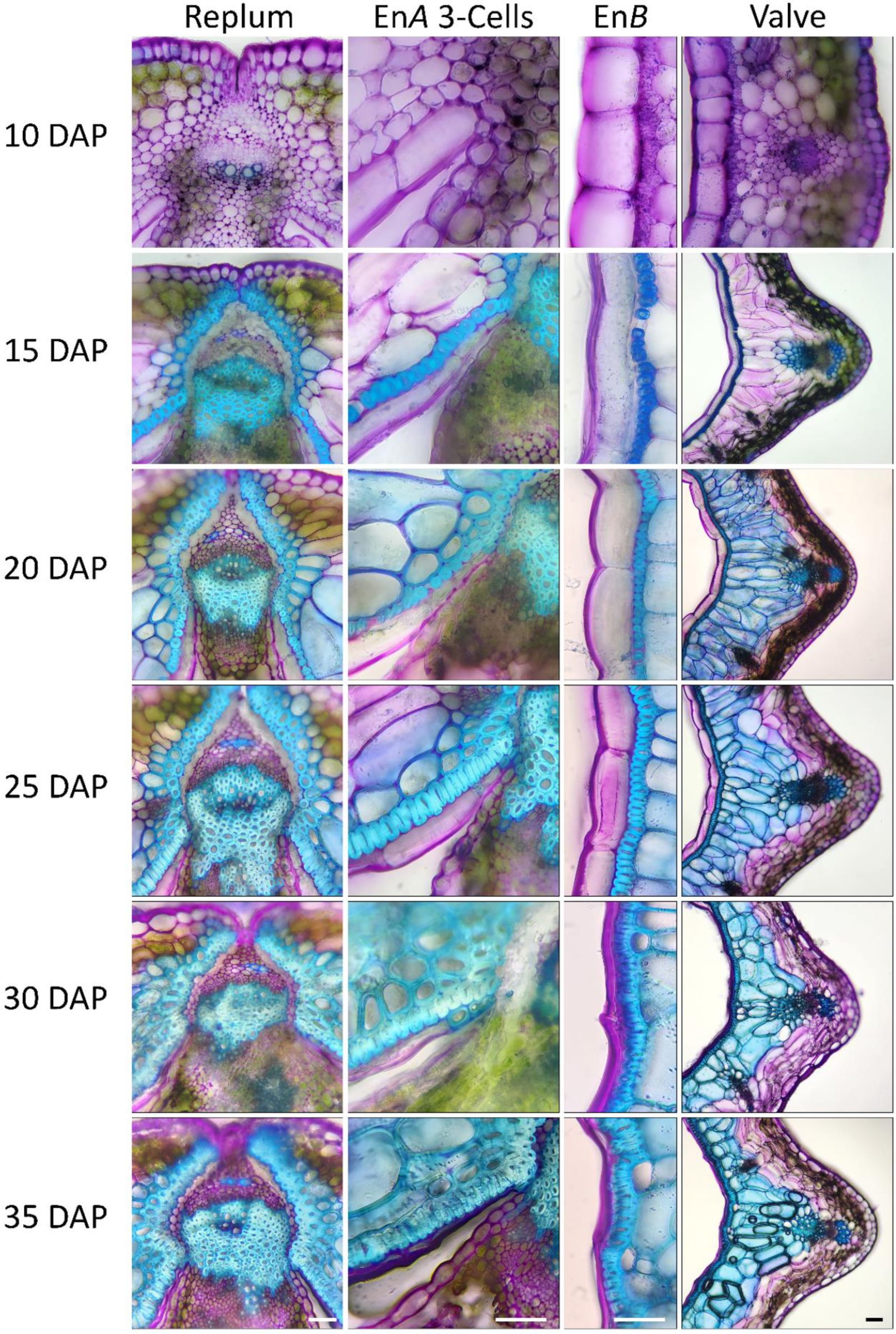
Spatial and temporal lignification of *Brassica nigra* siliques. Whole silique cross-sectional images 10, 15, 20, 25, 30, 35 days after pollination (DAP) depicting the replum, En*A* 3-Cells, En*B*, and valve spatial and temporal lignification patterning. Scale bars = 25 µm for each of the images in the panel.

